# SMoRe GloS: An efficient and flexible framework for inferring global sensitivity of agent-based model parameters

**DOI:** 10.1101/2024.09.18.613723

**Authors:** Daniel R. Bergman, Trachette Jackson, Harsh Vardhan Jain, Kerri-Ann Norton

## Abstract

Agent-based models (ABMs) have become essential tools for simulating complex biological, ecological, and social systems where emergent behaviors arise from the interactions among individual agents. Quantifying uncertainty through global sensitivity analysis is crucial for assessing the robustness and reliability of ABM predictions. However, most global sensitivity methods demand substantial computational resources, making them impractical for highly complex models. Here, we introduce SMoRe GloS (<u>S</u>urrogate <u>Mo</u>deling for <u>Re</u>capitulating <u>Glo</u>bal <u>S</u>ensitivity), a novel, computationally efficient method for performing global sensitivity analysis of ABMs. By leveraging explicitly formulated surrogate models, SMoRe GloS allows for comprehensive parameter space exploration and uncertainty quantification without sacrificing accuracy. We demonstrate our method’s flexibility by applying it to two biological ABMs: a simple 2D cell proliferation assay and a complex 3D vascular tumor growth model. Our results show that SMoRe GloS is compatible with simpler methods like the Morris one-at-a-time method, and more computationally intensive variance-based methods like eFAST. SMoRe GloS accurately recovered global sensitivity indices in each case while achieving substantial speedups, completing analyses in minutes. In contrast, direct implementation of eFAST amounted to several days of CPU time for the complex ABM. Remarkably, our method also estimates sensitivities for ABM parameters representing processes not explicitly included in the surrogate model, further enhancing its utility. By making global sensitivity analysis feasible for computationally expensive models, SMoRe GloS opens up new opportunities for uncertainty quantification in complex systems, allowing for more in depth exploration of model behavior, thereby increasing confidence in model predictions.

## 1 Introduction

Scientists today are generating abundant data and information as they seek to improve our comprehension of the world around us, revealing the inherent complexity characteristic of biological, biomedical, ecological, social, and other real-world systems. Agent-based models (ABMs) have emerged as a significant tool for understanding such complex systems, being particularly well-suited to capturing emergent phenomena^1–4^. ABMs are stochastic computational models that describe populations as individuals or agents, each with its own set of properties and behaviors that interact with their local environment to generate global phenomena. Such a formulation allows ABMs to capture connectivity and heterogeneity across multiple time, spatial, and structural scales^3,5^.

However, the use of ABMs presents significant challenges and drawbacks. For instance, the computational costs of solving ABMs escalate and become prohibitive when simulating millions of agents^5,6^. Furthermore, there is an absence of closed-form expressions linking ABM output with input parameters, making it hard to assess whether the results of ABMs are robust to parameter perturbations^7^. Moreover, as ABMs are increasingly applied to model highly complex biological and environmental systems, the number of input parameters grows, introducing greater uncertainty in parameter values. This uncertainty in model inputs will necessarily propagate to model outputs, raising questions about model accuracy and reliability.

Parameter sensitivity analysis is a common practical technique used to quantify uncertainty in model outputs as a function of uncertainty in the inputs, helping us better understand the limitations of the model^8^. This type of analysis identifies which input parameters – and, by extension, the biological, physical, or real-world processes they represent – are the most critical determinants of an output of interest^9^. Sensitivity analysis can be either local, assessing the effect of individual input parameters, or global, evaluating the combined influence of multiple parameters varied simultaneously across their full ranges^10^. For highly nonlinear models with a large number of estimated parameters, global sensitivity analysis is essential for drawing meaningful conclusions. Several methods have been developed for sensitivity analysis in parametric models, including variance-based methods, moment-independent techniques, Monte Carlo methods, and methods using spectral analysis (for recent reviews, see^11, 12^).

Simple global sensitivity analysis methods include one-at-a-time methods like the Morris method (MOAT), which is computationally efficient, having a cost scaling as ∼10 × the number of parameters^13^. However, MOAT provides only limited information and is best suited for factor prioritization or preliminary screening of model parameters. Additionally, MOAT cannot account for parameter interactions, which are often expected in nonlinear models, limiting its usefulness in more complex systems^9^. For more robust insights, variance-based methods such as the extended Fourier Amplitude Sensitivity Test (eFAST) or Sobol indices are generally preferred. These methods are capable of both factor prioritization and factor fixing, where the goal is to reduce uncertainty by identifying and fixing unimportant parameters. Additionally, these methods can account for interactions between parameters when computing model variance. However, these techniques come with a much higher computational cost, scaling as ∼10^3^× the number of parameters^9,14,15^. Regression-based methods, like Partial Rank Correlation Coefficient (PRCC), may be employed for factor mapping, which aims to identify important inputs within specific output domains. These methods also have high computational costs, lying somewhere between MOAT and eFAST^11,16^. Aside from MOAT, the computational expense of simulating complex models remains a major challenge when applying global sensitivity methods to ABMs. Long run times often render any meaningful sensitivity analysis of such models impractical^17^. As a result, sensitivity analysis of complex, computationally expensive ABMs is frequently omitted or only partially performed^7,18^.

One approach to addressing some of the aforementioned issues is to employ surrogate models, also known as metamodels or response surfaces. These are computationally less expensive models designed to approximate the dominant features of a complex model, here, the ABM^19^. Widely applied across various domains, surrogate models facilitate the exploration of ABM parameter spaces without incurring prohibitive computational costs^20–23^. Notably, surrogate model generation via Machine Learning, where the surrogate model does not have a closed form, is becoming increasingly popular^24^. However, such black box models have limited applicability in scenarios with limited training datasets or when extrapolating across broad and uncertain ABM parameter space where the a priori unknown ABM output could have high variability^25,26^. To mitigate these issues, we have proposed employing *explicitly formulated* surrogate models for approximating ABM behavior. Our approach has proven effective in parameterizing computationally complex ABMs with multi-dimensional data^5,6^. This work introduces a novel application of this technique to address the acute shortage of fast and accurate computational techniques for performing global sensitivity analysis of large-scale, complex ABMs.

Specifically, we develop a new, computationally efficient method, <u>S</u>urrogate <u>M</u>odeling for <u>R</u>ecapitulating <u>G</u>lobal <u>S</u>ensitivity (SMoRe GloS), that uses explicitly formulated surrogate models to infer the global sensitivity of input parameters in ABMs describing complex real-world systems. Our method is agnostic to any specific method for global sensitivity analysis and is easily adapted per user specification. To demonstrate our approach, we consider two spatio-temporally resolved ABMs representing biological processes: (1) an easy-to-simulate ABM representing a cell proliferation assay on a two-dimensional grid and (2) a more complex ABM of three-dimensional vascular tumor growth. SMoRe GloS computes the global sensitivity indices of ABM parameter sets in both instances using two techniques, namely, the computationally efficient MOAT Method and the computationally expensive but more versatile eFAST method. Remarkably, our method generates global sensitivity indices even for those ABM parameters that represent biological processes not explicitly included in the surrogate model formulation. We also compute sensitivity metrics directly in both instances and compare the results with our indirect method to validate our approach. Finally, we demonstrate the significant computational efficiency of SMoRe GloS compared to directly implementing methods like eFAST.

## 2 Methods

### 2.1 SMoRe GloS: <u>S</u>urrogate <u>M</u>odeling for <u>R</u>ecapitulating <u>G</u>lobal <u>S</u>ensitivity

Our new method for global analysis of computationally complex models, SMoRe GloS, is implemented in five steps: (1) Generate ABM output; (2) Formulate candidate surrogate models; (3) Select a surrogate model; (4) Infer relationship between surrogate model and ABM parameters; and (5) Use relationship between surrogate model and ABM parameters to infer global sensitivity of ABM parameters. These are described in further detail below.

We illustrate SMoRe GloS with two ABMs: one describing an *in vitro* cell proliferation assay that can be simulated easily and quickly; and one describing vascular tumor growth in 3-dimensions that is computationally complex and more expensive to simulate. These are described in further detail in subsections 2.3 and 2.4.

For convenience, we introduce the following notation. We will refer to the input ABM parameters to be included in the global sensitivity analysis as 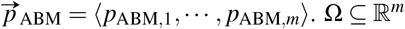, together with a probability distribution ρ, will denote the minimal sample space of these parameters. Parameters appearing in the surrogate model will be denoted 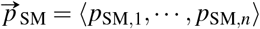. Finally, we will refer to surrogate model as SM.

#### Step 1: Generate ABM output

Sample ABM parameter values over Ω, making sure to include points along the boundary of Ω, together with some interior points. Aim for good coverage of Ω, bearing in mind the increased computational expense as more parameter values are selected. For this, choose any sampling method such as a regular grid, Latin Hypercube Sampling (LHS), random sampling, etc., considering each has advantages and disadvantages^27,28^. Next, generate ABM output at each sampled parameter vector, making sure to run multiple simulations in order to get meaningful averaged behavior.

In both our examples, we sampled ABM parameters on a regular grid, taking an average of *N* = 6 runs per sampled parameter vector.

#### Step 2: Formulate candidate surrogate models

Formulate (several) candidate SMs informed by the complex system being studied, the mechanisms encoded within the ABM, ABM output generated in Step 1, and most importantly, the output metric of interest in which we want to quantify the relative influence of each ABM parameter. More details on formulating explicit SMs are available here:^5,6^. Ideally, arrive at several candidate SMs.

For the *in vitro* cell proliferation ABM, our output metric of interest was total cell number at the end of the simulation. We therefore chose cell numbers in G1/S and G2/M phases of the cell cycle as the SM variables, and a system of two coupled ordinary differential equations (ODEs) describing their temporal evolution as the SM itself (see^6^ for more details). For the 3D vascular tumor growth ABM, our output metrics of interest were: (1) final tumor volume; (2) area under the tumor volume time-course; and (3) time to half-maximum tumor volume. These were chosen to illustrate various features and overall robustness of our method. Since ABM output was being integrated over space in all three instances, we once again used ODEs to formulate the SM, taking total cell number as the SM variable. Three candidate SMs were formulated in this case, namely, exponential growth, logistic growth and von Bertalanffy growth (see^5^ for more details). The SMs together with the corresponding ABMs are listed in subsections 2.3 and 2.4.

#### Step 3: Select a surrogate model

Select the best candidate from the various SMs formulated in Step 2 as follows. Considering each SM in turn, begin by fitting the SM to ABM output generated at each sampled ABM parameter vector (Step 1). In this process, make sure to collect information on <u>goodness-of-fit</u> of, and <u>uncertainty</u> in, the fitted SM parameters (discussed below). For the given SM, aggregate this information across all ABM output. Repeat this process for every candidate SM.

##### Goodness-of-fit criteria

Fit the SM to ABM output by maximum likelihood estimation (MLE)^29^, weighted least squares optimization^30^, or other method of parameter estimation. Record the quality of the fit.

In both our examples, we used weighted Residual Sum of Squares (RSS) to quantify goodness-of-fit.

##### Uncertainty in SM parameters

Quantify the uncertainty in SM parameters by computing confidence bounds when fitting the SM parameters to ABM output generated from each sampled ABM parameter vector. These confidence bounds will be used later, in Step 4. Several methods may be employed for uncertainty quantification (see for instance^19^).

Also quantify how well constrained SM parameters are by noting the span of their confidence bounds. For this, we propose a metric we call the **identifiability index**, which is defined as follows. If both upper and lower confidence bounds on an SM parameter are tightly-constrained when fitting to the ABM output generated at a sampled ABM parameter vector, the identifiability index is assigned a value of 2. Here, tightly-constrained parameters should have confidence bounds well within their physically or biologically relevant ranges. Parameters with one-sided confidence bounds, constrained only at one end, receive an identifiability index value of 1, while a score of 0 indicates an unconstrained parameter that may assume any value within its overall range. Thus, as the SM is fit in turn to all ABM output, a high frequency of 2’s will suggest an overall well-constrained SM parameter, whereas mostly 0’s will suggest unidentifiability of that parameter, possibly due to an over-parameterized SM.

In our examples, we used the profile likelihood approach^31–33^ to generate 95% confidence bounds on SM parameters. Identifiability indices were computed by graphing the likelihood curves obtained by profiling each fitted SM parameter. These cross the 95% confidence bound threshold never (a flat curve), once (an L-shaped curve), or twice (a U-shaped curve) times in the neighborhood of its best-fit value. The respective identifiability index values are 0, 1 or 2.

SM Selection: Select the best SM by considering both the goodness-of-fit and the identifiability index. The goal is to choose an SM that both minimizes RSS scores across ABM output, and has well-constrained SM parameters, as evidenced by a high frequency of 2’s in their identifiability indices. If selecting between SMs with different numbers of free parameters, model selection theory should be applied, for instance, by computing an Information Criterion^34^.

For the *in vitro* cell proliferation ABM, we did not need to perform model selection since we started with a single SM. For the 3D vascular tumor growth ABM, we reported the results of implementing SMoRe GloS with all three SMs, although a single SM emerged as the best overall candidate, based on our selection criterion outlined above. The Akaike Information Criterion (AIC) in Equation 1 below was used to aid in model selection.

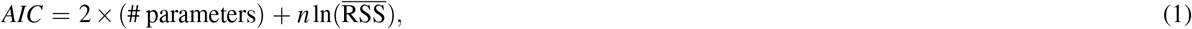

where 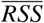 is the average RSS taken over *n* data points. Models with higher ΔAIC scores are less likely to explain the data. To compare between models, we computed a relative log-likelihood (*RLL*), defined as

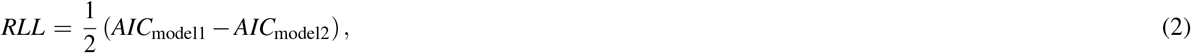

where a positive value of RLL indicates that model 2 is preferable to model 1.

#### Step 4: Infer relationship between SM and ABM parameters

Quantify the functional relationship between ABM parameters and SM parameters as follows. View each SM parameter as an unknown function – or hypersurface – of the ABM parameters. The (95%) confidence bounds on SM parameters inferred in Step 3 then correspond to discrete points on upper and lower (95%) confidence hypersurfaces ‘above’ the given ABM parameter vector, yielding a range of values for all SM parameters corresponding to each ABM parameter vector. These ranges are usually an interval for each SM parameter. The Cartesian product of these intervals – a hyperrectangle – defines the region of SM parameter space that best fits ABM output at that ABM parameter vector. These Cartesian products quantify the ‘stiff and sloppy’ nature of SM parameters^35^, providing information about the directions of SM parameter space that produce small (sloppy) or large (stiff) changes in model behavior. In particular, as the ABM parameter vector is varied, the deformations of these hyperrectangles give rise to variations in ‘stiffness and sloppiness’, which are used to determine ABM parameter sensitivities in Step 5. For more details on how to generate SM parameter hypersurfaces, refer to^5^.

#### Step 5: Use relationship between surrogate model and ABM parameters to infer global sensitivity of ABM parameters

Select an output metric of interest, say *f*, on the ABM and a method for computing the global sensitivity of *f* to changes in ABM parameters. *f* is a real-valued function on ABM parameter space, that is, *f* : Ω → ℝ. The global sensitivity, GS, is then a function of *f* and the probability distribution on ABM parameter space, *ρ*. Denote by GS (*f* (·); *ρ*) ∈ ℝ ^*m*^ the sensitivity of *f* to each of the *m* varied ABM parameters. The fundamental concept of SMoRe GloS is that an SM is used to estimate *f* in computing GS. Specifically, the value of *f* at an ABM parameter vector, 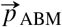, is approximated by sampling uniformly over the hyperrectangle in SM parameter space in Step 4 above. That is,

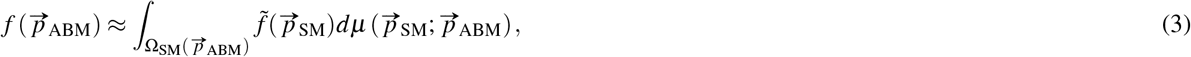

where 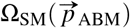 is the hyperrectangle in SM parameter space corresponding to 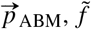 is the functional on SM parameter space to match *f*, and 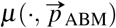 is the uniform probability distribution on 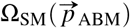. For notational simplicity, we will use *f* for 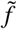 and *µ* for 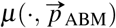 going forward. Putting this together with global sensitivity yields the following:

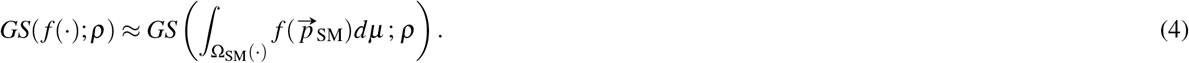

In our illustrative examples, we employ two methods for global sensitivity: the Morris Method and eFAST (see next section).

### 2.2 Global Sensitivity Analysis Methods

In this manuscript, we will illustrate how SMoRe GloS works using two global sensitivity methods: the Morris Method and eFAST (extended Fourier amplitude sensitivity test). The Morris Method is a one-step-at-a-time method that uses elementary effects (the effect of perturbing a single parameter) to compute a global sensitivity measure for each parameter^13,36^. This method has a low computational cost and its output is in the same units as that of the metric, making the sensitivity indices readily interpretable. Its main limitations are its inability to capture higher order interactions between model parameters and the fact that it does not yield a definitive boundary separating the important parameters from less influential ones. eFAST is a variance decomposition method that can efficiently handle models with nonlinear responses and complex interactions, and is model independent^14^. eFAST estimates the variance of the chosen model output, and the contribution of input parameters as well as their interactions to this variance. The algorithm then separates the output variance into the fraction of the variance that can be explained by variation in each input parameter. The result of this analysis is the main effect and total effect sensitivity indices.

### 2.3 Simple ABM of an *In Vitro* Cell Proliferation Assay

We consider the easy-to-simulate ABM presented in^6,37^, which describes a 2-dimensional on lattice birth-death-migration model of tumor cell proliferation. Briefly, cell division occurs as cell progress through four stages of the cell cycle in order: G1, S, G2, and M with transition rates, *ρ*_*G*1→*S*_, *ρ*_*S*→*G*2_, *ρ*_*G*2→*M*_, *ρ*_*M*→*G*1_, respectively. When a cell advances from M back to G1, it can proliferate into an unoccupied neighboring lattice site, provided the strength of contact inhibition on it is below a threshold *T*_*con*_. Otherwise, the cell returns to G1 without undergoing mitosis. Cells move to neighboring lattice sites at a constant migration rate, *s*, provided a randomly selected neighboring lattice site is unoccupied. If not, cells remain stationary. The growth culture is assumed to have a carrying capacity *K*_*A*_. For complete details on ABM formulation and simulation method, see^37^.

We infer global sensitivity of the seven ABM parameters mentioned above with respect to total cell number at the end of the simulation. These parameters are summarized in Table 1, and were varied across three values each, for a total of 3^7^ = 2187 ABM parameter vectors. At each of these, six replicates were simulated.

**Table 1.** List of ABM and surrogate model (SM) parameters

| Simple ABM Parameters |  | SM Parameters (equations (5)-(6)) |  |
| --- | --- | --- | --- |
| Parameter | Meaning | Parameter | Meaning |
| $K_A$ | Carrying capacity | $\lambda_C$ | G1/S → G2/M transition rate |
| $T_{con}$ | Contact inhibition | $\alpha_C$ | G2/M → G1/S transition rate |
| $s$ | Migration rate | $K_C$ | Carrying capacity |
| $\rho_{G1 \rightarrow S}$ | G1 → S transition | | |
| $\rho_{S \rightarrow G2}$ | S → G2 transition | | |
| $\rho_{G2 \rightarrow M}$ | G2 → M transition | | |
| $\rho_{M \rightarrow G1}$ | M → G1 transition | | |
| Complex ABM Parameters |  | SM Parameters (equations (7)-(9)) |  |
| Parameter | Meaning | Parameter | Meaning |
| $p_{div}$ | Progenitor cell proliferation rate | $\lambda$ | Exponential growth rate |
| $s_{div}$ | Stem cell proliferation rate | $r$ | Logistic growth rate |
| $r_{mig}$ | Tip cell migration rate | $K$ | Logistic carrying capacity |
| $p_{lim}$ | Progenitor cell division limit | $\alpha$ | vB growth rate |
| | | $\beta$ | vB death rate |
| | | $v$ | vB exponent |

Following^6^, an ODE formulation for the SM was chosen, with the numbers of cells in G1/S phase (*N*_1*S*_) and G2/M phase (*N*_2*M*_) as model variables. The following governing equations comprise the SM:

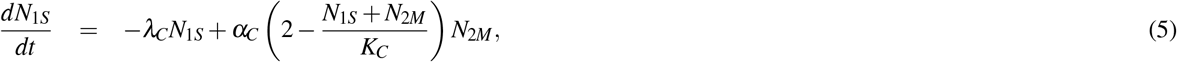

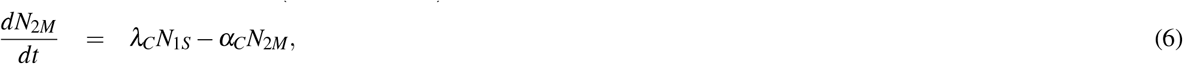

where λ_*C*_ is the rate of transition from G1/S to G2/M, *α*_*C*_ is the maximum rate of proliferation of cells in G2/M and *K*_*C*_ is the growth culture’s carrying capacity. For more details on how this SM was derived, see^6^. These parameters are summarized in Table 1.

### 2.4 Complex ABM of Vascular Tumor Growth in 3D

We consider the computationally complex model of vascular tumor growth in 3 dimensions presented in^38^. This on-lattice ABM consists of two modules that communicate with each other: a cancer cell module; and a vascular module.

The cancer cell module comprises cancer progenitor cells, which make up the bulk of the tumor, and cancer stem cells. The proliferation rate *p*_*div*_ of progenitor cells is greater than the proliferation rate *s*_*div*_ of cancer stem cells. Progenitor cells can divide a limited number of times, *p*_*lim*_, before they become senescent. On the other hand, cancer stem cells have limitless replicative potential. Progenitor cells reproduce symmetrically to produce two daughter progenitor cells, whereas cancer stem cells can reproduce asymmetrically or symmetrically, producing a progenitor daughter cell and a stem cell, or two stem cells. Both types of cancer cells migrate or proliferate only if there is space in an adjacent lattice site (Moore’s neighborhood). Both cell types are assumed to have a common migration rate, *mig*. A second factor governing the ability of a cancer cell to migrate or divide is its oxygen status, which could be normoxic (maximum migration and proliferation rates) or hypoxic (minimum migration and proliferation rates). This oxygen status is determined by the cell’s distance from a mature, blood-borne vessel.

The second module comprises endothelial cells and simulates angiogenesis: the formation of new blood vessels within the tumor. The tumor initially starts with a mature vasculature along its boundaries. As the tumor grows past the diffusion threshold of oxygen, the cancer cells become hypoxic. This triggers an ‘angiogenic-switch’ and cancer cells begin secreting Vascular Endothelial Growth Factor (VEGF), initiating angiogenesis. In response to this chemical stimulus, mature vessels near a hypoxic cancer cell can sprout, forming a new (non-mature) vessel. This sprout proliferates, extends, and migrates up the gradient of VEGF towards the nearest hypoxic cells until it anatamoses (fuses with) another sprout or with a nearby mature vessel. Once anastamosis occurs, the sprouts involved become blood-borne (mature) and nearby cancer cells become normoxic. We refer the reader to^5^ for complete details on this ABM and how to simulate this ABM.

We infer global sensitivity of the four ABM parameters mentioned above with respect to: (1) final tumor volume; (2) area under the tumor volume time-course; and (3) time to half-maximum tumor volume. These parameters are summarized in Table 1, and were varied across three values each, for a total of 3^4^ = 81 ABM parameter vectors. At each of these, six replicates were simulated.

An ODE formulation for the SM was chosen, with the total number of tumor cells (*N*) as the model variable. Three possible formulations were chosen for the SM, since each of these is a well-established model for tumor growth^39,40^:

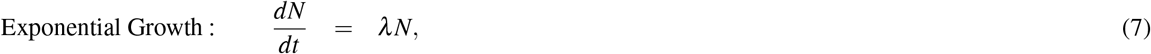

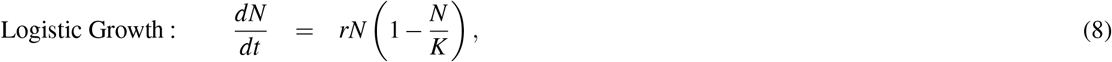

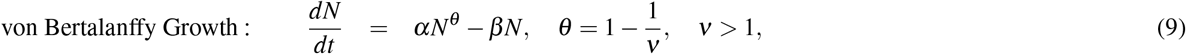

where λ is the exponential growth rate of tumor cells, *r* and *K* are the intrinsic growth rate and carrying capacity for the logistic model, respectively, and *α, β* and ν are the growth rate, death rate and exponent in the von Bertalanffy model, respectively. These parameters are summarized in Table 1.

## 3 Results

In this section, we demonstrate the accuracy of SMoRe GloS in computing the global sensitivity indices for ABM parameter sets through two distinct test cases. First, we explore an easy-to-simulate ABM that models an *in vitro* cell proliferation assay in two dimensions. Then, we apply our method to compute the global sensitivity of parameters in a more complex ABM that simulates three-dimensional vascular tumor growth.

### 3.1 Global Sensitivity of Parameters in ABM Representing Cell Proliferation Assay

We begin by generating output for the easy-to-run ABM of a two-dimensional cell proliferation assay, described in Section 2.3. Figure 2A presents a storyboard depicting a typical simulation at various time points, illustrating the spatial distribution and cell cycle phase distribution of cells from Day 0 to Day 3. Figure 2B shows time series data of cell numbers in G1/S and G2/M phases of the cell cycle from a typical ABM simulation, highlighting the accumulation of cells in G1/S as the total number of cells approaches the carrying capacity and the virtual cell culture exhausts available space. ABM parameters, together with the biological processes they regulate, are illustrated in Figure 2C. Parameters that represent spatial processes are highlighted in yellow and include the rate of cell movement, *s*, and the contact inhibition parameter, *T*_*con*_. We note that the surrogate model chosen for this ABM, specified in equations (5) and (6), is independent of local spatial considerations and, therefore, does not explicitly incorporate the processes represented by these parameters.

**Figure 1.**
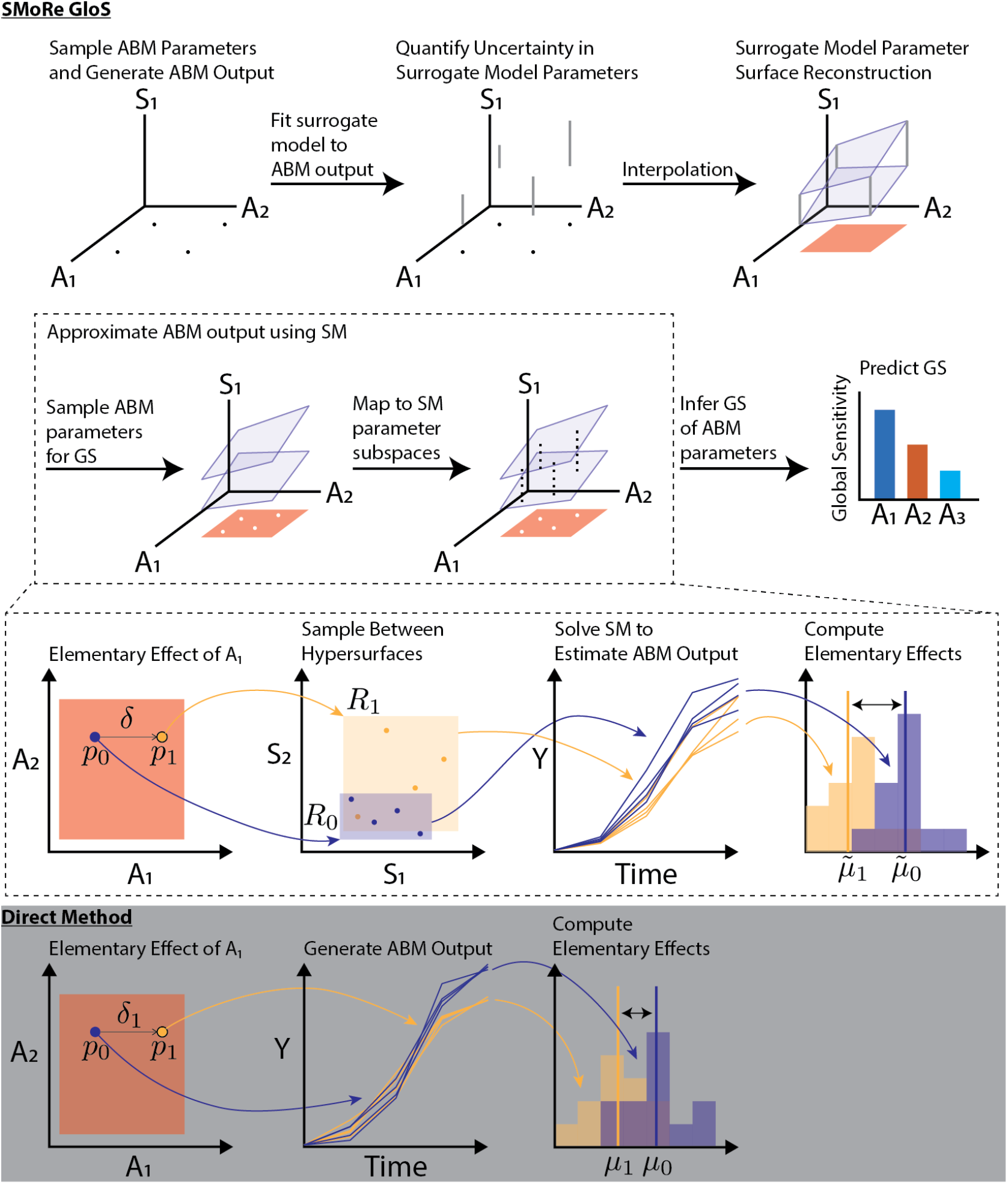
Schematic representation of the SMoRe GloS framework for sensitivity analysis of ABMs. For simplicity, two ABM parameters, *A*_1_ and *A*_2_, and one surrogate model (SM) parameter, *S*_1_, are depicted. The first row shows Steps 1-4 of SMoRe GloS, where *S*_1_ is constrained as a function of *A*_1_ and *A*_2_. The black dots represent sampled ABM parameters, the gray bars indicate uncertainty in *S*_1_ and the blue planes represent the reconstructed parameter surfaces for *S*_1_. The salmon region denotes the interior of the ABM parameter space, defined by the convex hull of the sampled points. The second row illustrates Step 5, where any global sensitivity method can be applied. The white dots represent points in ABM parameter space sampled for computing global sensitivity, and the dashed black lines show the corresponding ranges of *S*_1_. The third row illustrates the implementation of the MOAT method in this framework. Points *p*_0_ and *p*_1_ are examples of white dots from the second row that represent points in ABM parameter space used to compute an elementary effect in *A*_1_. These points correspond to regions *R*_0_ and *R*_1_ in SM parameter space. The time series curves are the trajectories sampled from these regions. The purple and yellow distributions denote the output metric of interest calculated from each trajectory. The elementary effect is approximated by the difference between the means of these distributions. The fourth row, with a dark background, illustrates the direct implementation of MOAT. Here, multiple ABM trajectories are generated at both *p*_0_ and *p*_1_, and the elementary effect of *A*_1_ is computed as before, using the difference between the means of the ABM output distributions.

**Figure 2.**
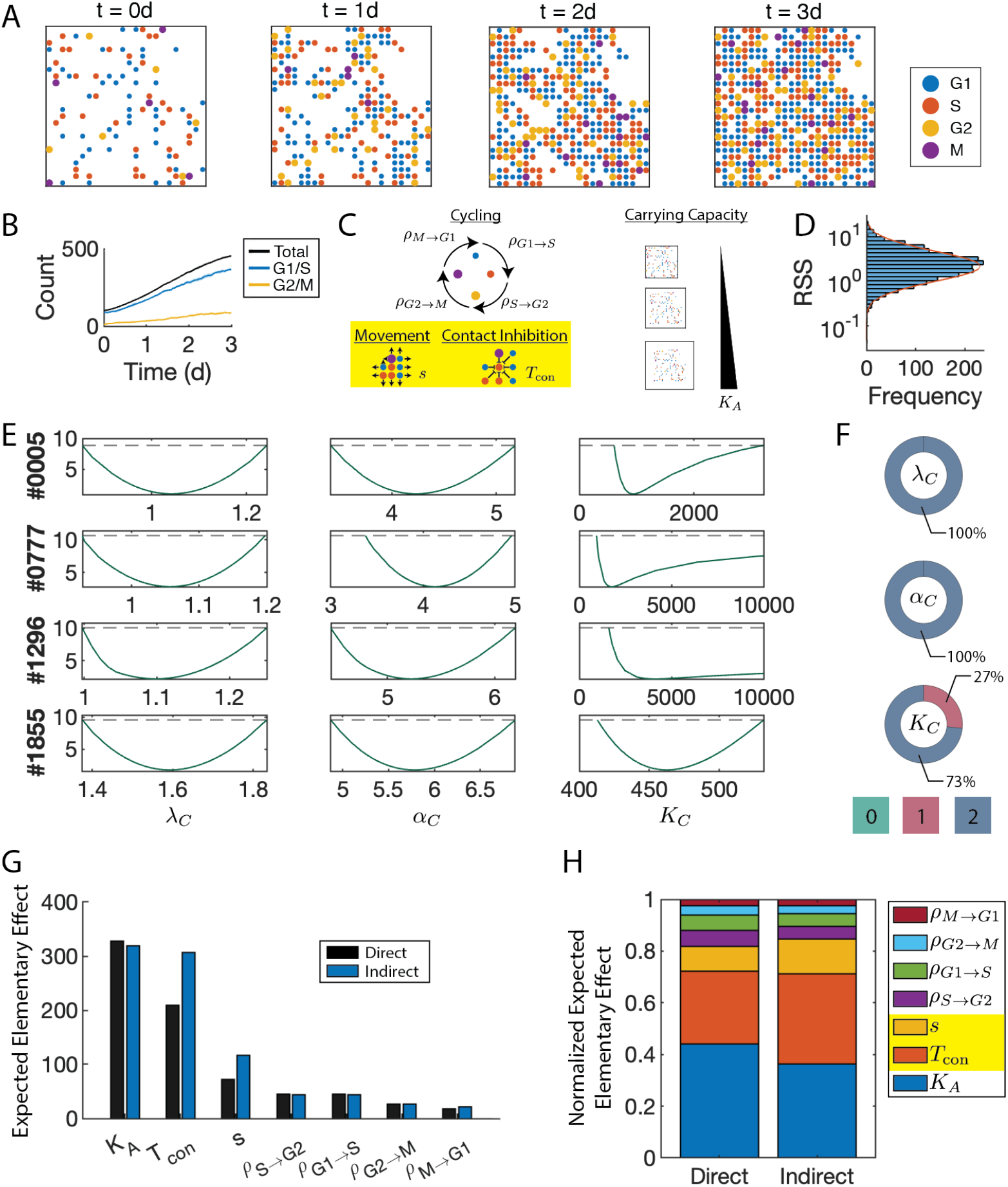
SMoRe GloS recapitulates global sensitivity of cell culture ABM. A) ABM storyboard showing cells by location and cell-cycle phase. B) Time series of the G1/S and G2/M cell-cycle phases. C) ABM parameters included in the sensitivity analysis. The yellow box highlights local spatial parameters that are not explicitly captured by the surrogate model (SM). D) RSS distribution of SM fits to all ABM parameter vectors. Orange line indicates the log-normal distribution that best fits this distribution. E) Profile likelihoods of SM parameters at four randomly selected ABM parameter vectors. F) Identifiability wheels of SM parameters where color indicates the identifiability index, and area the proportion of ABM parameter vectors for which the given SM parameter had that index. G) MOAT sensitivity analysis results using the ABM (Direct, black bars) and SMoRe GloS (Indirect, blue bars), ranked by decreasing sensitivity using the direct method. H) Normalized MOAT sensitivity values for each ABM parameter using the direct (left) and indirect (right) methods. Spatial parameters not not explicitly captured by the SM are highlighted in yellow.

#### 3.1.1 Surrogate Model Accurately Matches ABM Output with Minimal Uncertainty in Parameter Values

After selecting the best surrogate model, we fit it to the ABM output and calculate the residual sum of squares (RSS) to assess the goodness-of-fit (Step 3 of SMoRe GloS). The resulting distribution of RSS values is summarized in Figure 2D. The RSS values appear log-normally distributed with a very low mean (≈ 1), indicating an overall excellent fit quality. We also apply the profile-likelihood method, as described in Step 3 of SMoRe GloS, to quantify the uncertainty in surrogate model parameter estimates. Figure 2E shows sample profile likelihood curves for three surrogate parameters: *λ*_*C*_ (G1/S to G2/M transition rate), *α*_*C*_ (G2/M to G1/S transition rate), and *K*_*C*_ (carrying capacity), for four representative sets of ABM parameters. All likelihood profiles for *λ*_*C*_ and *α*_*C*_ are U-shaped and intersect the 95% confidence interval thresholds (dashed lines) twice. Consequently, their identifiability indices are 2 in each case. In contrast, the sample profile likelihoods for *K*_*C*_ can be L-shaped, intersecting the 95% confidence interval thresholds (dashed lines) only once. Thus, the identifiability index for *K*_*C*_ is 2 in the top and bottom cases shown, and 1 in the middle cases.

Aggregating across all ABM outputs, *λ*_*C*_ and *α*_*C*_ have consistently well-constrained upper and lower 95% bounds, with 100% of their identifiability indices having a value of 2 (Figure 2F, first two donuts). *K*_*C*_ exhibits some profiles identifiable from only one side, resulting in 73% of its indices being 2 and 27% being 1 (Figure 2F, bottom donut).

#### 3.1.2 SMoRe GloS Accurately Computes Global Sensitivity of 2D Cell Culture ABM Parameters, Including Those Not Explicitly Represented in the Surrogate Model

We next implement Steps 4 & 5 of SMoRe GloS to infer the global sensitivity of ABM parameters using two distinct methods: the Morris method and eFAST. In each case we also infer the sensitivity of ABM parameters directly using these methods, to evaluate the efficacy of SMoRe GloS. Fig 2G contrasts the global sensitivity of ABM parameters inferred directly (black bars) and indirectly using SMoRe GloS (blue bars) with the Morris method. Both approaches yield similar rankings for the importance of each parameter. The direct method suggests a higher sensitivity for carrying capacity compared to contact inhibition, though both were deemed highly sensitive by the indirect method as well. The direct and indirect methods are in excellent agreement on the insensitivity of transition rates between cell cycle phases and the intermediate sensitivity of cell migration rates. Fig 2H normalizes and stacks these sensitivities for clearer comparative visualization, reaffirming the ability of SMoRe GloS to accurately recapitulate the global sensitivity of ABM parameters using the Morris Method. Our method performs similarly well when using the eFAST method to infer global sensitivity of ABM parameters (see SI Figure S1).

These results showcase the capability of our method to infer the sensitivity of ABM parameters. Remarkably, this includes parameters representing local spatial processes (highlighted in yellow), such as cell movement and contact inhibition, which are beyond the scope of the surrogate model. It also extends to processes not explicitly included in the surrogate model, such as the transition rates from G1 to S and G2 to M.

### 3.2 Global Sensitivity of Parameters in ABM Representing 3-D Vascular Tumor Growth

Implementing Step 1 of SMoRe GloS for this case study, we generate output for a computationally complex ABM that models three-dimensional vascular tumor growth, as described in Section 2.4. Figure 3A presents a storyboard depicting a typical simulation at various time points, illustrating the growth of a tumor and its associated vasculature at various time points. ABM parameters, together with the biological processes they regulate, are depicted in Figure 3B. The rate of tip cell migration parameter *r*_mig_ represents a spatial process, and is highlighted in yellow. Following Step 2 of SMoRe GloS, three candidate surrogate models, specified in equations (7), (8) and (9)), are chosen for this ABM. It is important to note that these surrogate models are independent of spatial considerations and, therefore, do not explicitly incorporate the processes represented by *r*_mig_.

**Figure 3.**
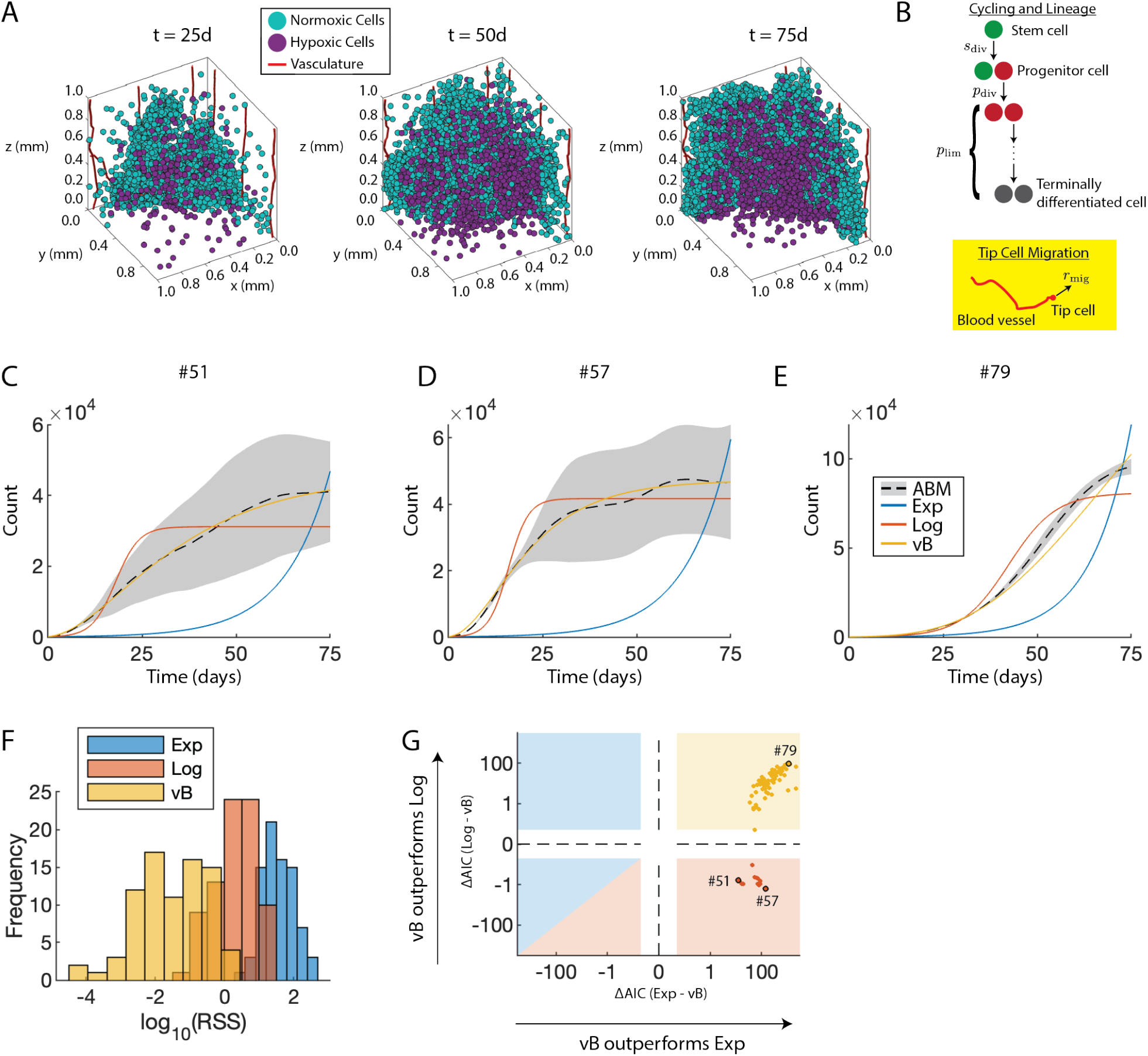
Surrogate Model (SM) selection for the 3D vascular tumor growth ABM. A) ABM storyboard showing vascular tumor growth. B) ABM parameters included in sensitivity analysis. The yellow box highlights local spatial parameters that are not explicitly captured by the SMs. C-E) Fits of the SMs to ABM output at three representative ABM parameter vectors. ABM parameter vectors were chosen based on the best fit to the exponential SM (C), logistic SM (D), and von Bertalanffy SM (E). F) Histograms of log10(RSS) values for each SM across all sampled ABM parameter vectors. G) Comparison of Akaike Information Criterion (AIC)-based relative log-likelihoods between the three SMs. Individual ABM parameter vectors are represented as darker colored dots. The x-axis shows the relative log-likelihood of the exponential model, and the y-axis shows the relative log-likelihood of the logistic model, both compared to the von Bertalanffy model. Positive (resp. negative) values indicate that von Bertalanffy is more (resp. less) likely than the alternative SM. The background is color-coded by the SM selected by AIC: yellow indicates preference for von Bertalanffy, red for logistic, and blue for exponential. The ABM parameter vectors corresponding to panels C), D), and E) are highlighted with black circles. Dashed lines indicate where the log scales change sign.

#### 3.2.1 Surrogate Model Selection for the Computationally Complex ABM is Guided by Goodness-of-fit and Identifiability Indices

Figures 3C-E show average cell number time courses (dashed lines), together with standard deviation (gray shaded area), from ABM simulations generated at three representative values of input parameters. Following Step 3 of SMoRe GloS, these figures also include fits of the three candidate SMs to the ABM output: exponential growth (blue curves, equation (7)); logistic growth (red curves, equation (8)); and von Bertalanffy growth (yellow curves, equation (9)). Visually, the von Bertalanffy model aligns more closely with the ABM output than the other two, while the exponential model performs the poorest. This observation is confirmed by the RSS distributions for the three models, shown in Figure 3F. The von Bertalanffy model provides a superior fit to the ABM output compared to the logistic and exponential models, as evidenced by a high frequency of low RSS values coupled with low variance. The exponential model yields the least accurate fits.

The above results are not surprising, given that the exponential model has one free parameter, the logistic model has two and the von Bertalanffy model has three. To facilitate model selection, the Akaike Information Criterion (AIC) is used to meaningfully compare the fits of the three surrogate models to ABM output, with results summarized in Figure 3G. This figure plots the relative log-likelihood of the von Bertalanffy model compared to the exponential (x-axis) and logistic (y-axis) models. The right half of the figure indicates when von Bertalanffy outperforms the exponential model, while the top half indicates when von Bertalanffy outperforms the logistic model. In particular, the yellow square represents all cases where von Bertalanffy is superior to both the exponential and logistic models (84% of cases). The red square and triangle represent all cases where logistic is superior to both von Bertalanffy and exponential models (16% of cases). In no instance is the exponential model superior to both von Bertalanffy and logistic models (blue square and triangle). The labeled dots correspond to the ABM parameters whose trajectories are shown in panels C-E.

Continuing to implement Step 3 of SMoRe GloS, we employ the profile-likelihood method to quantify uncertainty in the parameter values of all three surrogate models. Figures 4A, 4B, and 4C display representative profile likelihood curves for the exponential model (single parameter *λ*, blue curves), the logistic model (two parameters *r* and *K*, red curves), and the von Bertalanffy model (three parameters *α, ν*, and *β*, yellow curves), respectively. Figures 4D, 4E, and 4F show the corresponding identifiability index donut charts for these surrogate model parameters, aggregated over all ABM output. As can be seen, parameters in the exponential model (Figures 4A and 4D) and the logistic model (Figures 4B and 4E) have identifiability indices of 2 in almost all cases, suggesting these parameters are well constrained by the ABM output. In contrast, the identifiability indices for the von Bertalanffy model parameters *β* and *ν* are almost evenly distributed between 0’s and 1’s, and almost exclusively 1’s for *α*. This indicates that the von Bertalanffy model parameters are poorly constrained by the ABM output. Thus, even though the von Bertalanffy model provides the best quality of fit, as evidenced by low RSS values, the uncertainty in its parameter values is greatest.

**Figure 4.**
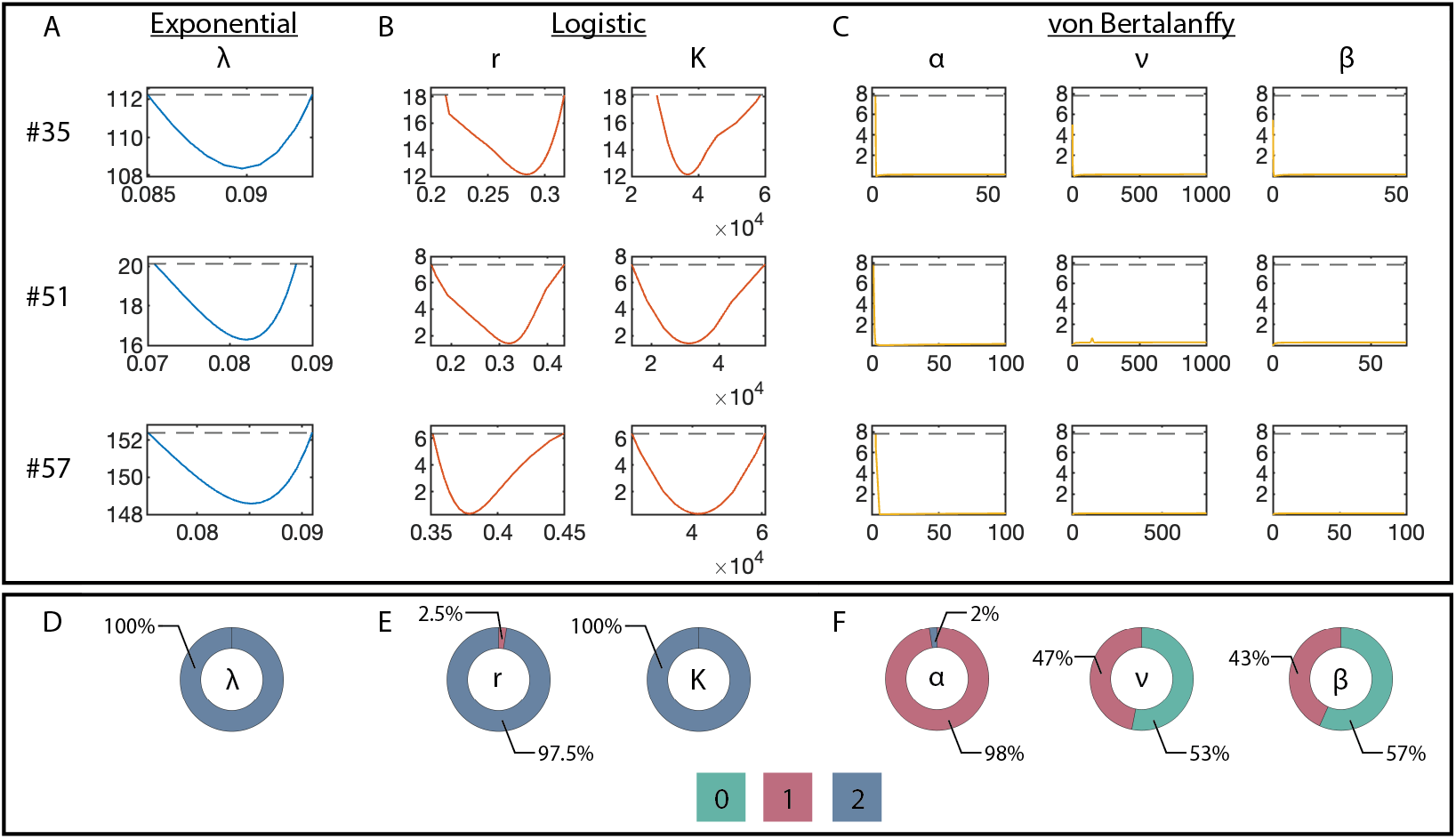
Comparison of the identifiability properties of the three surrogate models (SMs) for approximating the 3D vascular tumor growth ABM. A-C) Profile likelihoods for three representative ABM parameter vectors (rows) for each SM parameter (columns). D-F) Identifiability wheels of SM parameters where color indicates the identifiability index, and area the proportion of ABM parameter vectors for which the given SM parameter had that index. Each wheel is matched with the corresponding SM (columns A-C).

Considering these results, we expect the logistic model to perform best in the final step of SMoRe GloS due to its consistently good fits to ABM output and low uncertainty in parameter values. The exponential and von Bertalanffy only meet one of these criteria and are, therefore, not expected to yield optimal results.

#### 3.2.2 SMoRe GloS Accurately Computes the Global Sensitivity of ABM Parameters, with One Surrogate Model Emerging as the Best Choice

We now proceed to implement Steps 4 and 5 of SMoRe GloS to infer the global sensitivity of ABM parameters, employing two distinct methods: the Morris method and eFAST. We present below the results for MOAT. The results for eFAST are similar and can be found in SI Figure S2. To evaluate the efficacy of SMoRe GloS, we also directly infer the sensitivity of ABM parameters using these methods. For the global sensitivity analysis, we employ three distinct metrics to underscore the critical role of surrogate model selection in Step 3 of SMoRe GloS:

- final tumor size,
- area under the tumor volume time-course curve, and
- time to half-maximum tumor volume.

We selected these metrics based on their ability to capture different aspects of the data simulated by the ABM. Specifically, the final tumor size is independent of the dynamic properties of the tumor volume time-course, such as its shape and curvature. In contrast, both the area under the curve and the time to half-maximum volume are influenced to different degrees by these properties. These distinctions are illustrated in Figures 5A, 5D, and 5G. It is important to note that our choice of output metrics primarily aims to highlight the importance of surrogate model selection and does not necessarily reflect their biological relevance.

**Figure 5.**
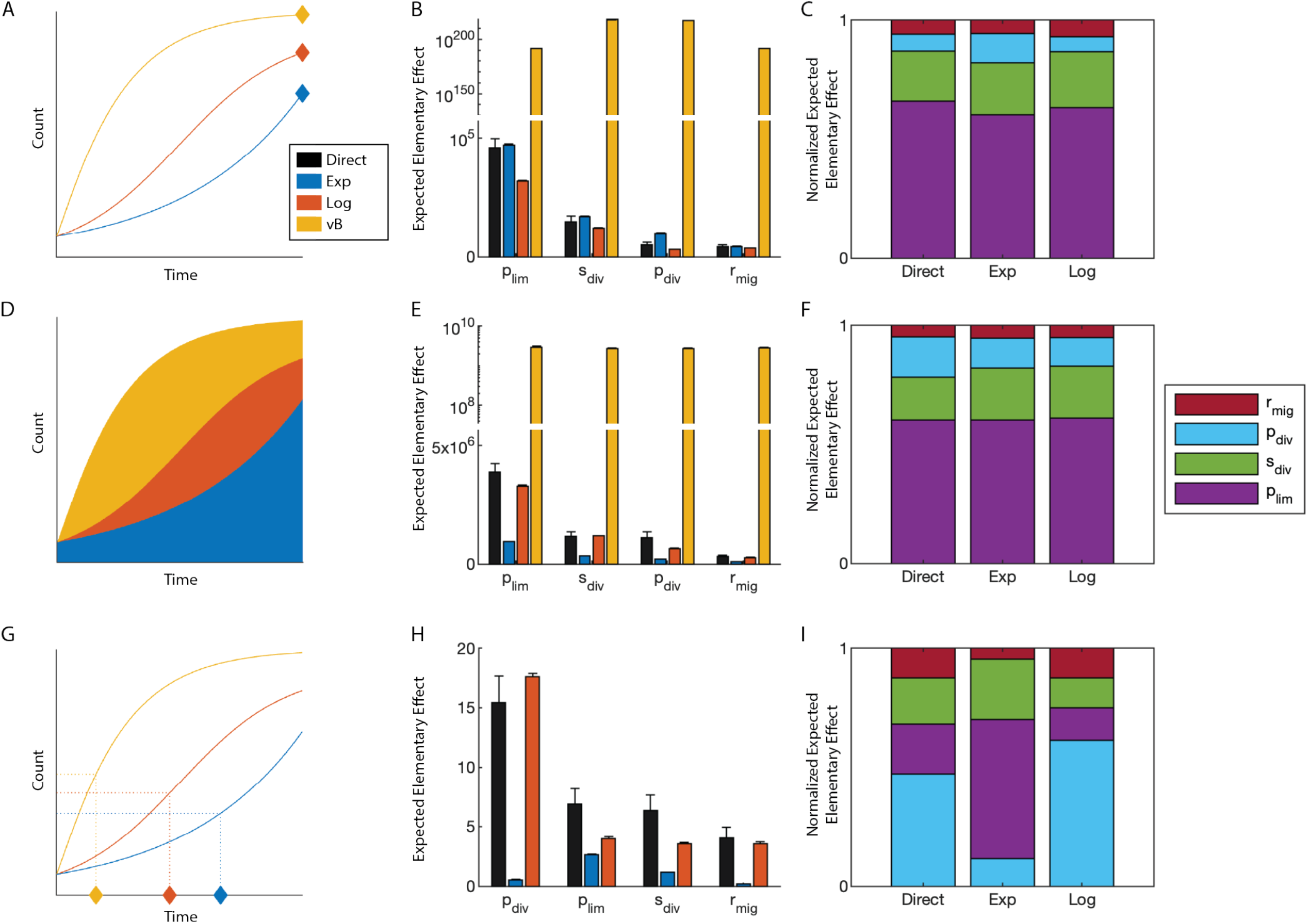
SMoRe GloS recapitulates global sensitivity of multiple output ABM metrics using the logistic surrogate model (SM). Each row uses a different output metric (left column) and shows the resulting sensitivity values (middle column) and their normalizations (right column). Colors in left and middle columns correspond to the SM as shown in the legend in A. Colors in the right column correspond to the ABM parameter as shown in the legend in F. A-C) Using final tumor size as the output metric. D-F) Using area under the curve as the output metric. G-I) Using time to half the maximum tumor volume as the output metric. Note the break in the y-axis scale in B and E.

Figures 5B, 5E, and 5H compare the global sensitivity of ABM parameters as inferred directly (black bars) and indirectly using SMoRe GloS with the Morris method across the three surrogate models (blue bars for the exponential model, red bars for the logistic model, and yellow bars for the von Bertalanffy model). Figures 5C, 5F, and 5I show the predicted relative importance of the ABM parameters for each metric by normalizing and stacking their sensitivities. The von Bertalanffy results are omitted from the normalization panels due poor unnormalized values.

##### Selecting a surrogate model solely based on goodness-of-fit to ABM output is insufficient for capturing global sensitivity

For all three global sensitivity metrics, the von Bertalanffy model – despite its superior fit to the ABM output – fails to adequately capture the sensitivity of the ABM parameters (Figures 5B and 5E, yellow bars). Notably, the time-to-half-maximum tumor volume results were so poor that they were not graphed (Figure 5H). This highlights the limitations of selecting a surrogate model based solely on goodness-of-fit fit to ABM output, without considering potential over-parameterization. Such an approach can severely compromise the effectiveness of the method.

##### Selecting a surrogate model solely based on minimizing uncertainty in its parameters is insufficient for capturing global sensitivity

The exponential and logistic models effectively predict the global sensitivities of ABM parameters with respect to final tumor size, as shown in Figure 5B (blue and red bars, respectively). The exponential model marginally outperforms the logistic model in capturing the sensitivity of the most significant parameter, while the logistic model excels in predicting the relative sensitivities of ABM parameters (Figure 5C).

Notably, the exponential model, which has the best identifiability indices, exhibits declining accuracy in calculating global sensitivity as the output metric becomes more reliant on the dynamic aspects of tumor growth. While it can accurately predict the order of importance of ABM parameters for the area under the tumor volume time-course curve (Figure 5E, blue bars), it fails to capture the true sensitivities of these parameters and completely fails when assessing the time to half maximum tumor volume (Figure 5H, blue bars). This is further evidenced by observing the predicted relative importance of ABM parameters (Figures 5C and 5F, second column versus first column).

##### Capturing global sensitivity accurately requires balancing good fits to ABM output with minimizing uncertainty in surrogate model Parameters

The logistic model consistently reproduces the sensitivities of ABM parameters across all evaluated metrics (Figures 5B, 5E and 5H, red bars, and Figures 5C, 5F and 5I, third column versus first column). These findings highlight the critical need to balance maximized goodness-of-fit with minimizing surrogate model parameter uncertainty when performing model selection in Step 3 of SMoRe GloS.

### 3.3 Computational efficiency of SMoRe GloS for Computing Global Sensitivity

The primary advantage of SMoRe GloS over directly computing global sensitivity with a complex model lies in its significant computational efficiency. Implementing the MOAT method directly with *d* parameters using a Latin Hypercube Sampling (LHS) of *k* points and *n*_*r*_ replicates at each point requires (*d* + 1) ×*k* × *n*_*r*_ ABM simulations. The *d* + 1 factor accounts for perturbing each LHS sample vector across all *d* parameter components. Typically, *k* values are recommended to range between 10 and 50^41^. For the 3D vascular tumor growth ABM, we varied *d* = 4 parameters using *k* = 15 LHS points, with *n*_*r*_ = 6 replicates, requiring 450 ABM simulations. Each simulation lasted, on average, 10 minutes, resulting in a total wall time of approximately 75 hours when run serially. In contrast, with SMoRe GloS, we started with the same (*d* + 1) × *k* = 75 ABM parameter points, but we drew 100 samples from the corresponding surrogate model (SM) parameter subspaces for each. This produced a total of 7,500 SM simulations. Since solving the SM has a negligible cost compared to interpolating the subspace and drawing samples, SMoRe GloS completed this task in under one minute (Figure 6A, blue line).

**Figure 6.**
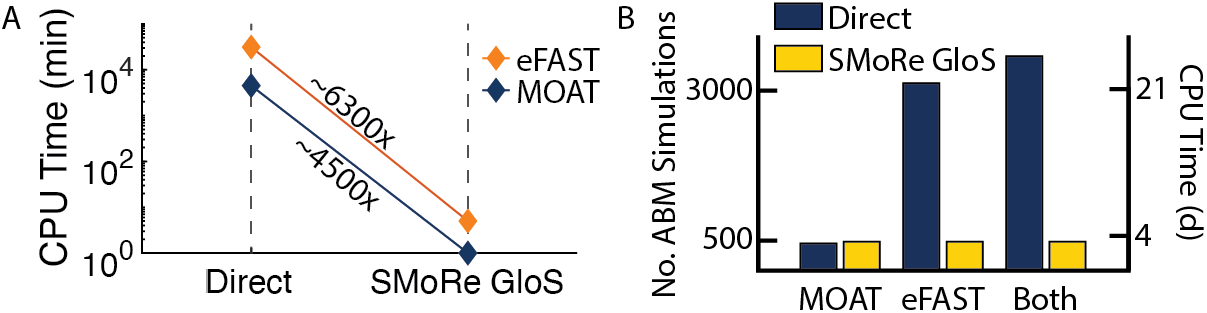
Comparison of ABM simulations and CPU time for computing global sensitivities using MOAT and eFAST in 3D vascular tumor growth ABM. A) Chart showing SMoRe GloS speedup (expressed as times faster) compared to direct implementation of global sensitivity analysis methods. The speedups exclude the setup time for the surrogate model. B) Number of ABM simulations and CPU time required to implement MOAT, eFAST, or both, either directly (blue bars) or with SMoRe GloS (yellow bars), including the setup time for the surrogate model. CPU time is based on assuming 1 ABM simulation takes 10 minutes.

For the more computationally intensive eFAST method, even more ABM simulations are required, further emphasizing the value of SMoRe GloS in improving computational efficiency. In our case, we applied eFAST to *d* = 4 parameters, with *N*_*r*_ = 2 replicates per parameter (corresponding to random phase shifts), and *N*_*s*_ = 65 samples per curve. The value *N*_*s*_ = 65 is the minimum recommended^14^. As with the MOAT method, we ran *n*_*r*_ = 6 replicates at each point to estimate the average ABM behavior. This led to a total of *d* × *N*_*r*_ × *N*_*s*_ × *n*_*r*_ = 3, 120 ABM simulations, which, if run serially, would require nearly 22 days of wall time. In contrast, SMoRe GloS once again demonstrated its computational superiority by completing the Efast analysis in under 5 minutes (Figure 6A, orange line).

SMoRe GloS does require an initial investment of computational resources for generating ABM output at sampled points in the ABM parameter space and profiling the SM against this output. For the vascular tumor growth ABM, we sampled *g* = 3 points in each of the *d* = 4 dimensions of parameter space, with *n*_*r*_ = 6 replicates at each point, resulting in a total of *g*^*d*^ ×*n*_*r*_ = 486 ABM simulations. While this number is comparable to the simulations required for directly computing MOAT sensitivities, it is significantly lower than what would be required for directly implementing eFAST. With just these 486 simulations, we were able to successfully recapitulate *both* MOAT and eFAST global sensitivity results. These are summarized in Figure 6B.

## 4 Discussion

In this paper, we introduce a novel method for inferring the global sensitivity of parameters in agent-based models (ABMs): Surrogate Modeling for Recapitulating Global Sensitivity (SMoRe GloS). This first-of-its-kind approach leverages explicitly formulated surrogate models to approximate ABM outputs, enabling a comprehensive exploration of parameter space that would otherwise be computationally prohibitive. Our findings demonstrate the potential of SMoRe GloS to significantly enhance the efficiency of global sensitivity analysis for ABMs, without compromising accuracy when applied judiciously.

One of the key strengths of SMoRe GloS is its combination of flexibility and adaptability. We demonstrated that our method performs consistently well with both eFAST and the Morris Method. By being agnostic to specific global sensitivity analysis techniques, SMoRe GloS offers greater compatibility across various sensitivity methods, with differing objectives like factor fixing, factor mapping and factor prioritization. This adaptability allows users to tailor the approach to their specific needs and preferences, which is particularly valuable given the wide range of applications for ABMs. Our successful application of SMoRe GloS to both, a two-dimensional cell proliferation assay, and a more complex three-dimensional vascular tumor growth model, highlights its broad utility.

SMoRe GloS offers significant computational efficiency compared to traditional approaches. For example, directly implementing the MOAT method for the 3D vascular tumor growth model required 450 ABM simulations, corresponding to ∼75 hours of CPU time, whereas SMoRe GloS achieved the same MOAT implementation in under 1 minute. The speedup was even more dramatic with eFAST, where direct implementation demanded 3,120 ABM simulations and 22 days of CPU time, while SMoRe GloS completed the task in under 5 minutes. Our results demonstrate that, even after accounting for the initial cost of setting up the surrogate model, SMoRe GloS provides substantial advantages in both speed and flexibility. This is particularly advantageous for more complex global sensitivity analysis tasks like factor mapping and prioritization, which are typically orders of magnitude more computationally expensive than simpler methods like MOAT, used for factor fixing.

We implemented SMoRe GloS with an on-grid parameter sampling, which scales exponentially with the dimensionality of the parameter space; this could be further optimized by employing Latin Hypercube Sampling (LHS), which scales linearly with parameter space dimensions. This would further reduce the computational cost of setting up the surrogate model. It is important to note that many complex models require hours per simulation, making direct global sensitivity analysis using methods like eFAST computationally prohibitive. However, SMoRe GloS makes such analyses feasible.

Another notable feature of SMoRe GloS is its ability to produce global sensitivity indices for ABM parameters that are not explicitly included in the surrogate model formulation. This feature enhances our method’s utility for complex models where certain biological or real-world processes are difficult to capture with computationally less expensive surrogate models. The implications are significant: we demonstrated that SMoRe GloS can accurately compute the sensitivity of spatial parameters that appear in an ABM, even when they are absent from a spatially-independent surrogate model.

One caveat of our approach is that the effectiveness of SMoRe GloS in accurately recovering the correct sensitivity indices of ABM parameters hinges on the choice of surrogate model. Ideally, one would aim to find a surrogate model that fits all ABM outputs near perfectly, with parameters that are fully identifiable – that is, determined with minimal uncertainty – across all outputs. However, this may be unattainable in practice because improvements in the fit quality frequently come at the cost of introducing additional parameters that may diminish their identifiability properties. To address this, we advocate for a balanced approach to surrogate model selection, guided by both goodness-of-fit to ABM output and the identifiability properties of surrogate model parameters. Specifically, the focus during surrogate model selection should be on ensuring it faithfully reproduces the ABM output with minimal uncertainty. Developing a mechanistic surrogate model that aligns with the underlying mechanisms coded in the ABM could be a promising strategy. The particular output metrics of interest, for which we wish to determine the sensitivities of ABM parameters, should be considered after selecting a robust surrogate model. Since a well-constrained surrogate model will be broadly applicable, it can effectively assess a variety of output metrics, making our approach particularly valuable given the unpredictable nature of exploratory modeling.

There are several promising avenues for further developing and extending SMoRe GloS. One potential direction under active consideration is to establish a ranking system for ABM parameters based on their influence on surrogate model parameters. This information could then be integrated with a sensitivity analysis of the surrogate model parameters to produce a global sensitivity ranking for the ABM parameters. Such an approach might eliminate the need to reconstruct surrogate model parameter hypersurfaces, thereby increasing our method’s efficiency. Additionally, as previously discussed, obtaining a well-constrained surrogate model that faithfully reproduces the ABM outputs of interest is crucial. To this end, we are currently exploring the use of machine learning and equation learning algorithms to further enhance our results. These approaches could lead to more robust and accurate surrogate models, ultimately broadening the applicability and efficiency of SMoRe GloS in various complex biological and real-world systems.

## Acknowledgements

The authors acknowledge generous support from the Institute for Computational and Experimental Research in Mathematics (ICERM) through their Collaborate@ICERM program, the American Institute of Mathematics (AIM) through their AIM SQuarREs program and the Mathematisches Forschungsinstitut Oberwolfach (MFO) through their Researchers in Pairs (RiP) program. This work was supported by NIH/NCI U01CA243075 (T.J.) and by NSF 2324818 (T.J, H.V.J. and K.-A.N.).

## Author contributions statement

All authors conceived the original idea and developed the mathematical methods. D.B. wrote the code and performed the numerical simulations. All authors analyzed the output. All authors wrote and reviewed the manuscript.

## References

1. Badham, J. et al. Developing agent-based models of complex health behaviour. Heal. & Place 54, 170–177 (2018).

2. Bianchi, F. & Squazzoni, F. Agent-based models in sociology. Wiley Interdiscip. Rev. Comput. Stat. 7, 284–306 (2015).

3. Bonabeau, E. Agent-based modeling: Methods and techniques for simulating human systems. Proc. Natl. Acad. Sci. 99, 7280–7287 (2002).

4. West, J., Robertson-Tessi, M. & Anderson, A. R. Agent-based methods facilitate integrative science in cancer. Trends Cell Biol. 33, 300–311 (2023).

5. Jain, H. V., Norton, K.-A., Prado, B. B. & Jackson, T. L. Smore pars: A novel methodology for bridging modeling modalities and experimental data applied to 3d vascular tumor growth. Front. Mol. Biosci. 9, 1056461 (2022).

6. Bergman, D. R., Norton, K.-A., Jain, H. V. & Jackson, T. Connecting agent-based models with high-dimensional parameter spaces to multidimensional data using smore pars: A surrogate modeling approach. Bull. Math. Biol. 86, 1–28 (2024).

7. Borgonovo, E., Pangallo, M., Rivkin, J., Rizzo, L. & Siggelkow, N. Sensitivity analysis of agent-based models: a new protocol. Comput. Math. Organ. Theory 28, 52–94 (2022).

8. Saltelli, A. Sensitivity analysis for importance assessment. Risk Analysis 22, 579–590 (2002).

9. Saltelli, A. et al. Global sensitivity analysis: the primer, chap. 1, 1–51 (John Wiley & Sons, 2008).

10. Zhou, X., Lin, H. & Lin, H. Encyclopedia of GIS, chap. Global Sensitivity Analysis, 408–409 (Springer US, Boston, MA, 2008).

11. Iooss, B. & Lemaître, P. A Review on Global Sensitivity Analysis Methods, 101–122 (Springer US, Boston, MA, 2015).

12. Borgonovo, E. & Plischke, E. Sensitivity analysis: A review of recent advances. Eur. J. Oper. Res. 248, 869–887 (2016).

13. Morris, M. D. Factorial sampling plans for preliminary computational experiments. Technometrics 33, 161–174 (1991).

14. Marino, S., Hogue, I. B., Ray, C. J. & Kirschner, D. E. A methodology for performing global uncertainty and sensitivity analysis in systems biology. J. Theor. Biol. 254, 178–196 (2008).

15. Saltelli, A., Tarantola, S. & Chan, K.-S. A quantitative model-independent method for global sensitivity analysis of model output. Technometrics 41, 39–56 (1999).

16. Pianosi, F. et al. Sensitivity analysis of environmental models: A systematic review with practical workflow. Environ. Model. & Softw. 79, 214–232 (2016).

17. Sheikholeslami, R., Razavi, S., Gupta, H. V., Becker, W. & Haghnegahdar, A. Global sensitivity analysis for high-dimensional problems: How to objectively group factors and measure robustness and convergence while reducing computational cost. Environ. modelling & software 111, 282–299 (2019).

18. Thiele, J. C., Kurth, W. & Grimm, V. Facilitating parameter estimation and sensitivity analysis of agent-based models: A cookbook using netlogo and r. J. Artif. Soc. Soc. Simul. 17, 11 (2014).

19. Smith, R. C. Uncertainty quantification: theory, implementation, and applications (SIAM, 2013).

20. Dosi, G., Pereira, M. C. & Virgillito, M. E. On the robustness of the fat-tailed distribution of firm growth rates: a global sensitivity analysis. J. Econ. Interact. Coord. 13, 173–193 (2018).

21. Palar, P. S., Liem, R. P., Zuhal, L. R. & Shimoyama, K. On the use of surrogate models in engineering design optimization and exploration: The key issues. In Proceedings of the genetic and evolutionary computation conference companion, 1592–1602 (2019).

22. Schultz, M. G. et al. Can deep learning beat numerical weather prediction? Philos. Transactions Royal Soc. A 379, 20200097 (2021).

23. Vlahogianni, E. I. Optimization of traffic forecasting: Intelligent surrogate modeling. Transp. Res. Part C: Emerg. Technol. 55, 14–23 (2015).

24. Ten Broeke, G., Van Voorn, G., Ligtenberg, A. & Molenaar, J. The use of surrogate models to analyse agent-based models. J. Artif. Soc. Soc. Simul. 24 (2021).

25. Brigato, L. & Iocchi, L. A close look at deep learning with small data. In 2020 25th International Conference on Pattern Recognition (ICPR), 2490–2497 (2021).

26. Kasim, M. F. et al. Building high accuracy emulators for scientific simulations with deep neural architecture search. Mach. Learn. Sci. Technol. 3, 015013 (2021).

27. Renardy, M., Joslyn, L. R., Millar, J. A. & Kirschner, D. E. To sobol or not to sobol? the effects of sampling schemes in systems biology applications. Math. Biosci. 337, 108593 (2021).

28. Urban, N. M. & Fricker, T. E. A comparison of latin hypercube and grid ensemble designs for the multivariate emulation of an earth system model. Comput. & Geosci. 36, 746–755 (2010).

29. Millar, R. B. Maximum likelihood estimation and inference: with examples in R, SAS and ADMB (John Wiley & Sons, 2011).

30. Cohen, A. & Migliorati, G. Optimal weighted least-squares methods. The SMAI J. Comput. Math. 3, 181–203 (2017).

31. Venzon, D. & Moolgavkar, S. A method for computing profile-likelihood-based confidence intervals. J. Royal Stat. Soc. Ser. C (Applied Stat. 37, 87–94 (1988).

32. Eisenberg, M. C. & Hayashi, M. A. Determining identifiable parameter combinations using subset profiling. Math. Biosci. 116–126 (2014).

33. Eisenberg, M. C. & Jain, H. V. A confidence building exercise in data and identifiability: Modeling cancer chemotherapy as a case study. J. Theor. Biol. 431, 63–78 (2017).

34. Anderson, D. & Burnham, K. Model selection and multi-model inference. Second. NY: Sprimger-Verlag 63, 10 (2004).

35. Gutenkunst, R. N. et al. Universally sloppy parameter sensitivities in systems biology models. PLoS Comput. Biol. 3, e189 (2007).

36. Bergman, D. R., Karikomi, M. K., Yu, M., Nie, Q. & MacLean, A. L. Modeling the effects of emt-immune dynamics on carcinoma disease progression. Commun. Biol. 4, 983 (2021).

37. Bergman, D. & Jackson, T. L. Phenotype switching in a global method for agent-based models of biological tissue. Plos one 18, e0281672 (2023).

38. Norton, K.-A., Jin, K. & Popel, A. S. Modeling triple-negative breast cancer heterogeneity: Effects of stromal macrophages, fibroblasts and tumor vasculature. J. Theor. Biol. 452, 56–68 (2018).

39. Ghaffari Laleh, N. et al. Classical mathematical models for prediction of response to chemotherapy and immunotherapy. PLoS Comput. Biol. 18, e1009822 (2022).

40. Sarapata, E. A. & De Pillis, L. A comparison and catalog of intrinsic tumor growth models. Bull. Math. Biol. 76, 2010–2024 (2014).

41. Campolongo, F., Cariboni, J. & Saltelli, A. An effective screening design for sensitivity analysis of large models. Environ. modelling & software 22, 1509–1518 (2007).

