## Supplementary Information for "SMoRe GloS: An efficient and flexible framework for inferring global sensitivity of agent-based model parameters"

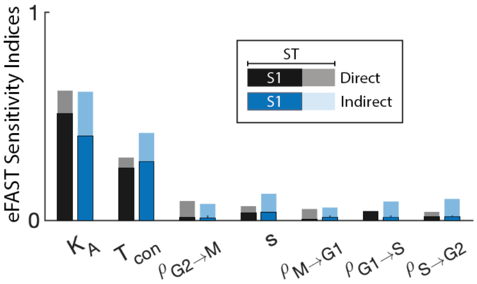


**Figure S1.** SMoRe GloS recapitulates eFAST global sensitivity of final tumor volume using the surrogate model (SM). Compare with Figure 2. Solid color, black-bordered bars represent first-order sensitivity indices. Transparent bars represent total-order indices. Note: these are not stacked bar plots; the total-order index is given by the height of the transparent bar, not the difference with the height of the first-order index bar.

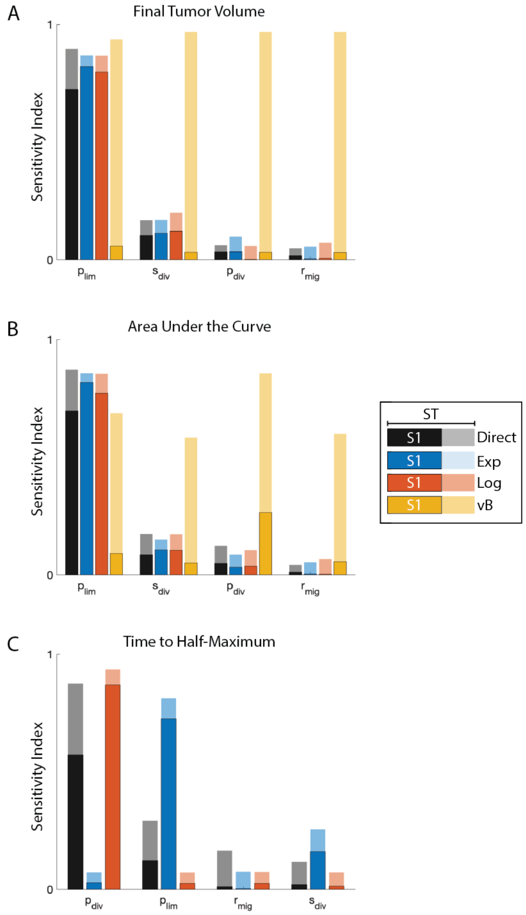


**Figure S2.** SMoRe GloS recapitulates eFAST global sensitivity of multiple output ABM metrics using the logistic SM. Compare with Figure 5. Solid color, black-bordered bars represent first-order sensitivity indices. Transparent bars represent total-order indices. Note: these are not stacked bar plots; the total-order index is given by the height of the transparent bar, not the difference with the height of the first-order index bar.
